# Synergistic population encoding and precise coordinated variability across interlaminar ensembles in the early visual system

**DOI:** 10.1101/812859

**Authors:** Daniel J Denman, R Clay Reid

## Abstract

Sensory stimuli are represented by the joint activity of large populations of neurons across the mammalian cortex. Information in such responses is limited by trial-to-trial variability. Because that variability is not independent between neurons, it has the potential to improve or degrade the amount of sensory information in the population response. How visual information scales with population size remains an open empirical question. Here, we use *Neuropixels* to simultaneously record tens to hundreds of single neurons in primary visual cortex (V1) and lateral geniculate nucleus (LGN) of mice and estimate population information. We found a mix of synergistic and redundant coding: synergy predominated in small populations (2-12 cells) before giving way to redundancy. The shared variability of this coding regime included global shared spike count variability at longer timescales, layer specific shared spike count variability at finer timescales, and shared variability in spike timing (jitter) that linked ensembles that span layers. Such ensembles defined by their shared variability carry more information. Our results suggest fine time scale stimulus encoding may be distributed across physically overlapping but distinct ensembles in V1.

## INTRODUCTION

The cortex represents information through the spiking activity of many neurons, distributed across both time and the neural population. In sensory and other neural systems, information can be extracted from many aspects of such responses. Historically, the variance in spiking rate of individual cells relative to the stimulus time is used (i.e., rate-based feature encoding, Adrian & Zotterman, 1926). Decades of work and substantial evidence indicates that visual information can be captured from such rate-based feature codes^2,3^, yet several lines of evidence suggest information may also be extracted from absolute spike times in the visual^4,5^, and other^6,7^ systems. The stimulus information in spike timing or spike rates depends on both the temporal precision and the shared variability of spikes within the sensory responses.

The temporal precision of spike times relative to stimulus presentations in single neurons responses has been measured across the early visual system in anesthetized^8–10^ some awake animals^11^. In the anesthetized visual system, individual neurons can be extremely precise, down to single milliseconds^12,13^. Such temporal precision has been hypothesized to be useful for spike-timing dependent plasticity^14,15^, exploiting dendritic non-linearities *in vitro*^16^ and *in vivo*^17^, enhancing transmission efficacy between areas^18–20^, activity routing^21–23^, and/or for timing-based codes^24,25^.

Yet, across trials both the timing and number of spikes is variable and limits information. What is the source of this variance? It is sometimes assumed that the process underlying spike time irregularity is inherently stochastic^26,27^, or subject to intrinsic noise in neural systems such as thermal noise ^28^, conductance noise^29^, synaptic release noise^30^, or could be reflecting chaotic dynamics^31^. Alternatively, variance could reflect variance in precisely timed inputs that are not under experimental control. Could the visual responses of single neurons be more precise if measured relative each other, or to an internal frame of reference of potential use to the system? If and how latency variability is correlated across cells in the awake central visual system is not known^9^.

This structure of correlated variability within a population, whether in the spike count or spike timing domain, profoundly impacts how stimulus information scales with the population size, whether in the latency domain as described above or in better-studied spike count domain^32–38^. Because state-of-the-art *in vivo* imaging approaches for recording populations of single neurons lack the temporal resolution and sensitivity to resolve individual spike times, the question of how sensory information (as represented in any potential aspect of spike trains, counts or times) scales with population size has remained in the domain of model-based approaches.

Our contribution here is an empirical assessment of how correlations in variability and stimulus information, change across scales: timescales down to single milliseconds, and as number of neurons scales up to hundreds of simultaneously recorded visual neurons. We find that, despite considerable trial-to-trial variability in the timing of individual neuron responses, populations synergistically encode stimulus information, including at ultrafine (< 10 msec) timescales. To account for such ultrafine-timescale information in apparently variable responses, we extend noise correlation analysis to temporal and population domains, showing that useful relative spike time information is maintained via strong, population-wide correlations in spike timing variability (i.e. jitter). This observation of strong correlated jitter and of usable additional stimulus information in population relative spike times supports the notion that the mammalian visual system can use dynamics distributed across layers and areas to encode stimulus information. We further find that maximal population information and correlated jitter is distributed across (not within) layers, suggesting that population encoding may be better be thought of as distributed and not as strictly hierarchical^39^.

## RESULTS

To study stimulus information and the coordination of spiking activity within populations in the early visual system, we recorded from populations of single neurons in visual cortex (V1) and lateral geniculate nucleus (LGN) of awake mice using high-density electrophysiology (*Neuropixels* probes^40^; Figure 1A-C) and repeatedly presented the same visual stimuli. The total number of single units (classified as fast-spiking (FS) or regular-spiking (RS) in cortex, Fig S1A-E) recorded simultaneously in each recording varied from 31-174 (Figure 1C). For more details on recordings, see Methods and Figure S1 and S2.

**Figure 1.**
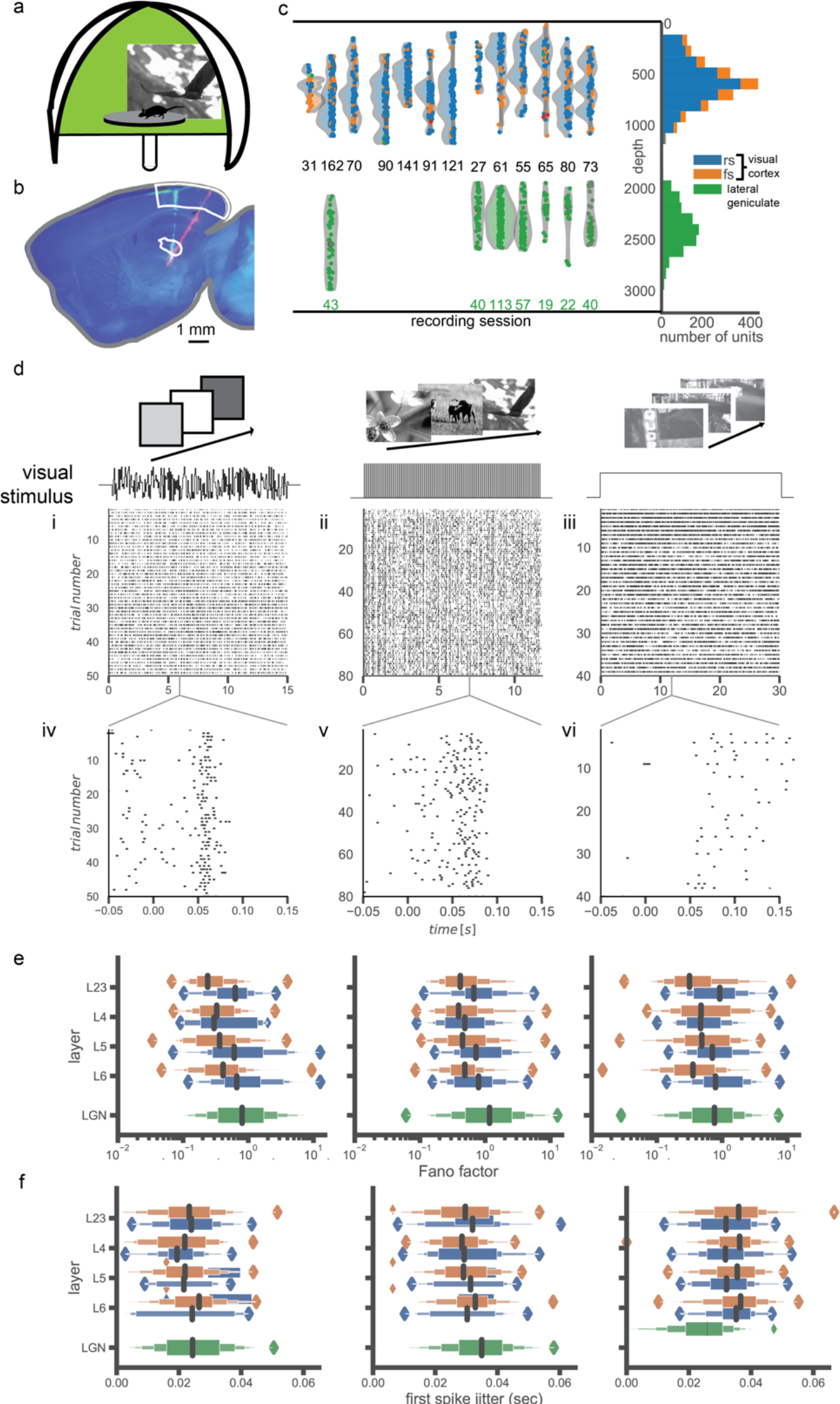
Recording from large populations of single cells across dLGN and V1 simultaneously. **a**., Top left: cartoon depicting the immersive visual stimulation system, with mouse in the center of a spherical enclosure at 30 cm viewing distance and an example naturalistic image from the stimulus set. **b.**, histological reconstruction of dual tracks through dLGN and V1. Sagittal slice, with DAPI in blue, one track from DiI on probe (anterior, pseudo-colored green), and one track in DiO (posterior, pseudo-colored red). **c.**, Distribution of units across V1 depth (left), for both regular- (blue) and fast-spiking (orange) units; dLGN units are shown for recordings that also penetrated dlGN in green. **d.**, Example responses to each class of visual stimulus. For each stimulus the temporal profile is shown above an example raster from the same example cell; for flicker, the luminance steps are shown, for naturalistc images, the timing of transitions from image-to-image, for the naturalistic movie, a block of continuous presentation is indicated. *i-iii.*, raster for the full duration of each stimulus type across all trials. *iv-vi.*, 200 msec zoom of response events for each stimulus type. **e.**, Fano factor of spike counts by area, layer, and cell type for each stimulus type **f.**, first spike jitter (standard deviation of first spike time) by area, layer, and cell type for each stimulus type

### Trial-to-trial variability in awake mice

We began by assessing how variable the spiking responses of single visual neurons are to repeated presentations of stimuli. Previous work has measured such variability in spiking - in other species, in other brain states (particularly anesthetized^10,12,41^, and at slower timescales. We sought to assess spiking variability across a range of timescales in the awake state.

We repeatedly presented a series of fixed-sequence (i.e., repeated) stimuli: a spatially-uniform luminance “flicker” stimulus (Figure 1Di), which varied in a random but fixed sequence; 118 naturalistic images presented in the same sequence (Figure 1Dii); and a repeated 30 second naturalistic movie clip (Figure 1Diii,^42^). Each stimulus was repeated 40-100 times, depending on the recording session.

In the spike count domain, our data recapitulate the findings from anesthetized animals that variance scales with mean^43,44^, can become sub-Poisson especially at higher firing rates^45^ (Fig S3), and that spike count variance is dependent on the stimulus type^33^ (Fig S3) in both V1 and LGN. Mean Fano factor across all cells in LGN was 1.11 - 1.99, depending on the stimulus and 0.66 - 0.91 in V1 (Figure 1E, Fig S3), slightly lower than has previously been reported for Ca^2+^ imaging in awake mice (1.39, ref. ^46^).

In the temporal domain, we measured the variability in spike timing, or jitter. Because of the known dependence of temporal precision of visual responses on the timescale of the visual stimulus^47^, we chose stimuli that covered a range of timescales: luminance flicker at 20 Hz, images presented at 10 Hz, and the “naturalistic” movie with variable but generally slower temporal frequency content (weighted average of time-frequency spectrogram: 3.4 Hz, Fig S4). While responses to each stimulus type were selective and grossly repeatable, in that each unit responded with visible “events” around distinct features of each stimulus (vertical stripes in individual raster plots, Figure 1Div-vi), trial-to-trial spike time varied considerably (Figure 1Div-vii).

We first examined the variability in the first spike time across trials (Fig S5). As expected, mean absolute latency was lowest in LGN (44.8 ± 0.7 msec), followed by V1 layer 4 (48.9 ± 0.9 msec) and subsequent V1 layers (layer 5: 49.9 ± 0.8 msec; layer 6: 52.0 ± 0.9 msec; layer 2/3: 53.0 ± 0.8 msec). The standard deviation of this first spike time across trials (“first spike jitter”) was lower in layer 4 (27.7 ± 0.5 msec, Figure 1F) than any cortical layer (layer 2/3: 29.2 ± 0.4 msec; layer 5: 28.7 ± 0.5 msec; layer 6: 30.0 ± 0.5 msec, Figure 1F). First spike jitter in LGN (29.7 ± 0.3 msec) was higher than layer 4, but not significantly different than any other cortical layer (p > 0.1, Welch’s t-test). Responses were more precise (less jitter) with increasing temporal frequency content in the stimulus (ref ^47^; 23.3 ± 0.003 msec for flicker, 31.1 ± 0.003 msec for naturalistic image stimuli, and 33.4 ± 0.003 msec for natural movies). The jitter in the median spike of each response was 3 - 4 msec smaller than the first spike jitter, but the relative median spike jitter across areas and stimuli was consistent with first spike jitter (Fig S5). Notably, the responses reported here in awake mice (LGN jitter: 24.6 msec) are considerably less precise than those reported in anesthetized cat (1-8 msec,^10,12^ and anesthetized mouse (8.8 msec^48^).

In summary, in awake mice both the spike count and spike timing of single neurons are significantly variable, even in response to stimuli that evoke high firing rate responses.

### Synergistic and redundant stimulus encoding in visual populations

Such trial-to-trial variability limits single neuron information coding capacity, and is thought to also limit population coding capacity. To first quantify the total stimulus information available in single neuron spike trains we estimated, across time scales, the maximum information rate in bits/sec. For single cells, we used the “direct” method ^12,49^. For each cell, we computed the total stimulus information by subtracting the response noise entropy (H_noise_) from an estimate of the total response entropy (H_total_), each of these H extrapolated to infinite word length from estimates of entropy at several word lengths (Figure 2A, see Methods). We found that the amount of information about the stimulus increased with increasing temporal resolution (decreasing bin size, Figure 2B), reaching 4.3 +/− 0.1 bits/s for LGN and 2.4 +/− 0.13 bits/sec for V1 (Figure 2C). LGN cells had the highest median information (2.77 bits/sec), and 93.5% of LGN units contained appreciable stimulus information (> 0.5 bits/sec). Regular-spiking cells had more information than FS cells in each layer (L2/3: RS= 2.37 ± 0.29, FS= 1.14 ± 0.09, p<0.001; L4: RS= 3.30 ± 0.70, FS: 1.60 ± 0.17, p = 0.03; L5: RS=2.87 ± 0.39, FS= 1.46 ± 0.10, p< 0.001; L6: RS= 2.66 ± 0.38, FS= 1.32 ± 0.09, p< 0.001). This was true for each stimulus type (Sup Fig 6). While the relative information rates between LGN and V1 - and bin sizes - were consistent with anesthetized cat^10^, the absolute amounts of stimulus information are nearly an order of magnitude lower in these awake mouse recordings.

**Figure 2.**
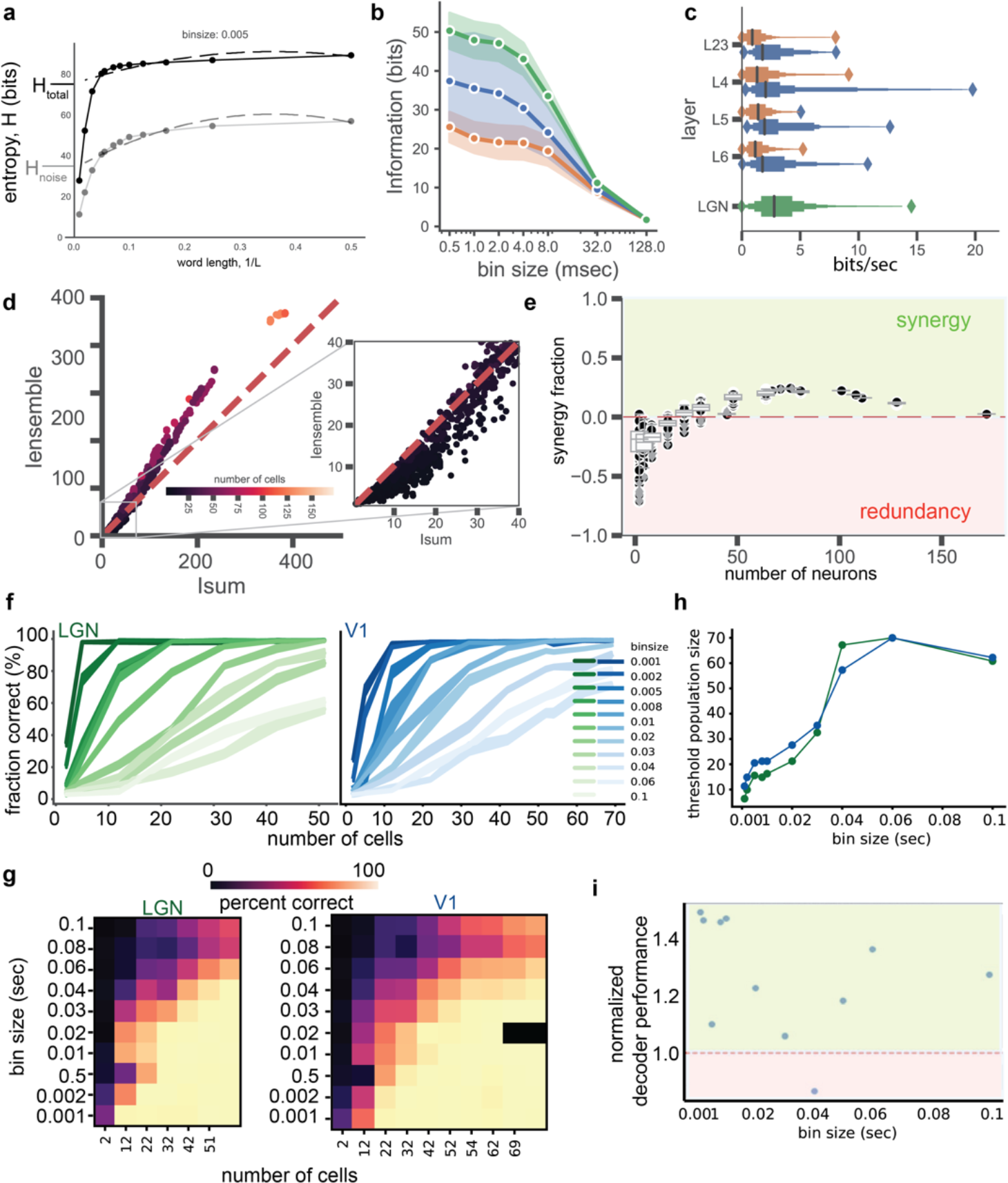
Information and stimulus decoding in dLGN-V1 populations. **a**., Estimation of entropy and information for individual cells. **b.**, stimulus (flicker) information as a function of bin size for V1 regular-spiking (blue), V1 fast-spiking, (orange) and dLGN (green) cells. Mean (points) and standard deviation (shading) shown. **c.**, letter value box plots of stimulus (flicker) information rate (bits/sec), broken down by layer. **d.**,I_sum_ and I_ensemble_ for all populations measured. For populations smaller than the total number recorded, subsets were repeatedly chosen and I_sum_ and I_ensemble_ recalculated. **e.**, synergy fraction,(I_ensemble_ − I_sum_) / I_ensemble_ for each population in (**e**). Box plots with 95% confidence intervals in grey. Outliers in black. Red and green shading highlight redundancy and synergy, respectively. **f.**, Performance of linear support vector machine classifier in decoding responses of increasing population size, at various temporal resolutions (shading), from an example recording including only dLGN or only V1 (regular and fast-spiking cells). **g.**, Performance of linear support vector machine classifier across all recordings, for dLGN and V1 populations. **h.**,The number of cells in either dLGN (green) or V1 (blue) needed to achieve a 70% correct criterion, as a function of bin size. Not the large dips in population size below both 40 and 5 msec. **i.**, Normalized decoder performance in decoding simultaneously recorded responses vs responses of the same cells from non-simultaneous trials. Red and green shading highlight redundancy and synergy, respectively.

Previous work has suggested that pairs of neurons in LGN and V1 can be synergistic, in which the simultaneous activity carries more information than the sum of the individual neurons^8,50,51^, redundant with less than the sum of individual neurons^52^, independent^52^, or both synergistic and redundant^53^. We addressed this in awake mice, and extended to ask how stimulus information grows with increasing population size. To do so we estimated stimulus information in ensembles of simultaneously recorded neurons using previously introduced methods^53^; this information is termed ensemble information, or I_ensemble_. We directly compared I_ensemble_ to the summed information (I_sum_) of each cell in the population used to compute I_ensemble_. I_ensemble_ = I_sum_ indicates independent coding, I_ensemble_ > I_sum_ indicates synergistic coding, and I_ensemble_ < I_sum_ indicates redundant coding. We observed both synergistic and redundant population (Figure 2D-F, Fig S6), depending on the population size.

Pairs of neurons in awake mouse dLGN and V1 recapitulate what has been observed for pairs of neurons in anesthetized macaques^52,53^. We observed weakly redundant (22.0%) coding between pairs of cells in both dLGN and within V1 (Figure 2D,E, Fig S6C). Growing the number of neurons beyond pairs did not significantly change the amount of redundancy in populations, up to 8 cells (Figure 2E, p > 0.05, Fig S6C). Beyond approximately 12 cells, increasingly large populations shifted toward synergy (where synergy is defined as more information in the combined response than expected from the information in individual responses, Figure 2E). The synergy began to return towards net independence after ~60 cells. The peak synergy fraction values we observed for the total population of each recording was 19.6 ± 2.9% (from 62 ± 23 cells, mean ± s.d.). Overall, our result indicate that, on average, weak pairwise redundancy co-exists within synergistic small populations. Beyond these small synergistic populations, adding more neurons begins to accumulate more redundancy.

To further understand the structure of population-level information, and because information theory based measures of the importance of correlated activity can be difficult to interpret^54^, we trained and tested several statistical decoders on our population responses to naturalistic images. The performance of these decoders in classifying which image generated a population response can be treated as a proxy for the available encoded stimulus information. We explored several decoders, choosing to use a linear support vector machine (SVM) because it performs at least as well as other decoders ^55^ and because the resulting decoder weights indicate the relative contribution of each element to decoding.

The accuracy of SVM decoding of image identity from population responses to held-out image presentations increased with more neurons, saturating before reaching the full population size (Figure 2F). Performance increased faster when using spike counts in smaller bins, continuing to improve down to the smallest bin size possible (1 msec, Figure 2F,G). The interaction of spike count bin size and number of neurons on decoding accuracy is shown in Figure 2G. Both finer temporal resolution and more neurons increase performance. To summarize this interaction we measured the populations required to achieve a standard psychophysics performance criterion, 70% correct. This summary showed that the number of neurons the decoder needed to successfully “do the task” of image classification dropped precipitously when smaller bin sizes were used, with a further dip when bin sizes reached down to single spike resolution (< 2 msec; Figure 2H).

To ask if the decoder was sensitive to within-trial population temporal structure (as suggested by the observed population synergy in Figure 2E), we compared the performance of standard and “shuffled trial” decoding. For “shuffled trial” decoding, the decoder was trained on simultaneously recorded population responses, but tested using pseudo-trials synthesized from a randomly selected trial for each neuron in a recording. This shuffling of the test responses made the decoder less accurate (Figure 2I), indicating that useful trial-to-trial information is present at all timescales. The enhancement in stimulus information within simultaneously recorded populations versus the shuffled responses of those same neurons was strongest with finer than 20 millisecond resolution (1 – 10 msec, Fig 2J).

In summary, in awake mice the visual stimulus information available in LGN and V1 increased with population sizes from 8 to 40 neurons, and at temporal resolutions < 40 msec. Because the structure of how variability is shared across neurons is known play a profound role in how information scales with population size^35^, we next explicitly measured this structure of correlated variability, pairwise and population, within and across areas in both the spike count and spike time domains.

### Correlated spike count variability across timescales

We began by measuring the correlated variability in spike count, or noise correlation (r_sc_) at a resolution similar that used in the literature (100 msec). Mean r_sc_ was 0.27 ± 0.001 in LGN and 0.13 ± 0.001 in V1 (Figure 3A) and was weakly and positively correlated with signal correlation (Fig S7A). For flicker, r_sc_ was 0.09 ± 0.001 between simultaneously recorded cells across V1 and LGN, slightly lower than V1 (0.10 ± 0.001, p = 0.0004, Welch’s t-test) and LGN (0.19 ± 0.001, p < 1e^−16^, Welch’s t-test), indicating at least one source of shared variability that correlates V1 pairs independently of LGN inputs to V1. The dependence of r_sc_ on stimulus type was opposite for V1 and LGN, with V1 r_sc_ higher for the flicker stimulus and LGN r_sc_ higher for the naturalistic movie stimulus (Fig S7B).

**Figure 3.**
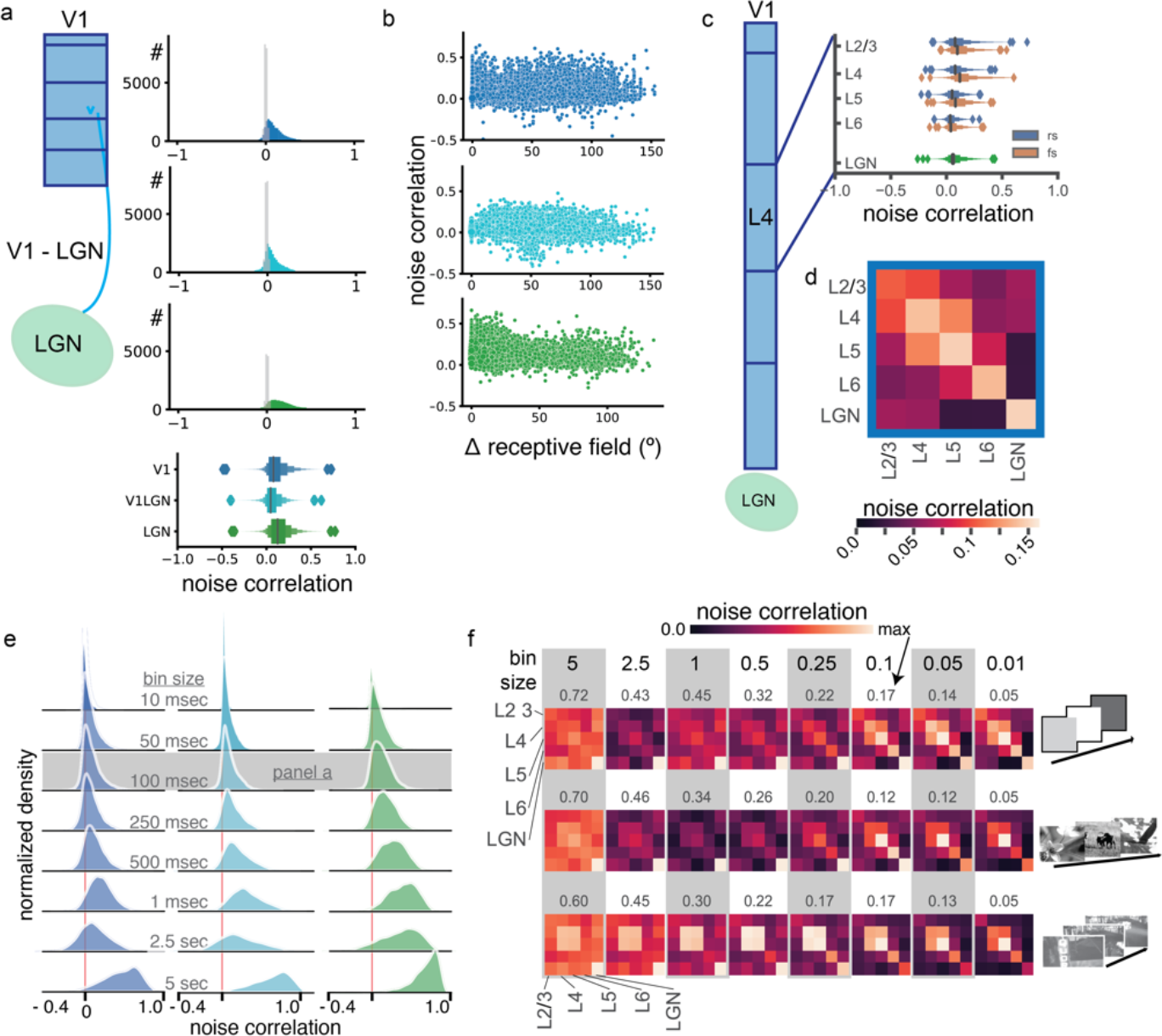
Correlated spike count variability across timescales, layers, and areas in dLGN-V1 populations. **a.,** spike count correlated variability (noise correlation) for V1-V1 pairs (dark blue, top), dLGN-V1 pairs (light blue, middle),and_dLGN-dLGN pairs (green, bottom). On each a null distribution from the same responses with trials shuffled for each cell is plotted in grey. **b.**, noise correlation as a function of receptive field offset for V1-V1 pairs (dark blue, top),_dLGN-V1 pairs (light blue, middle),and_dLGN-dLGN pairs (green, bottom). **c.**, letter value box plots of spike count correlated variability (noise correlation) for V1 layer 4 pairs. All pairs contain at least one cell from layer 4. **d.**, spike count correlated variability (noise correlation) for V1 layer and LGN pair combinations. **e.**, spike count correlated variability (noise correlation) for V1-V1 pairs (dark blue, left),_dLGN-V1 pairs (light blue, middle),and_dLGN-dLGN pairs (green, right) across timescales. Each row shows the spike count correlations for spike counts within the binsize indicated in grey. Red line indicates 0. **f.**, spike count noise correlation across layers and areas, for each temporal resolution (bin size, columns) and each stimulus (rows). For clarity, each color map is normalized to its maximum, which is shown above each matrix.

Expanding the analyses beyond the 100 msec bin size, we found spike count correlated variability depended strongly on timescale^32,56^. For <100 msec bin sizes we saw tighter distributions of r_sc_ with lower means (naturalistic images, 50 msec; LGN-LGN: 0.08 ± 0.001, LGN-V1: 0.03 ± 0.001, V1-V1: 0.07 ± 0.001, Figure 3C). At longer timescales r_sc_ was much higher: (naturalistic images, 50 msec; LGN-LGN: 0.42 ± 0.002, LGN-V1: 0.19 ± 0.002, V1-V1: 0.17 ± 0.001, Figure 3C). At most timescales, intra-V1 layer and intra-LGN r_sc_ was higher than across class r_sc_ (Figure 3D). However, the manner in which pairwise r_sc_ depended on timescale was not uniform across layers or across areas.

At longer timescales, we found that correlated variability reflected a small number of global influences, consistent with recent reports^57,58^. In V1, all layers are correlated with all other layers (Figure 3D, leftmost columns of 1, 2.5, and 5 sec bins); within LGN correlated variability was particularly high at the longest timescale measured, 5 seconds (0.67 ± 0.002), and the across-area (dLGN-V1) r_sc_ was also strengthened when measured at longer timescales (0.51 ± 0.002).

At shorter timescales (< 150 msec), we found overall lower correlation in spike counts (Figure 3D) and more across-layer structure. Higher temporal resolution noise correlations revealed variability within and across layers that differentiated from a global phenomenon (Figure 3F). Most notably, layers 4 and 5 became particularly correlated in variability at fast time scales; layer 2/3 became more isolated from the other cortical layers.

Thus, correlated spike count variability depends on timescale and pair location within the dLGN-V1 network. The differences in noise correlation across timescales underscore that interpreting noise correlations at a single timescale may mask different network phenomenon, even from networks that have the same underlying anatomical connectivity.

### Correlated spike time variability (jitter)

In addition to spike counts, we assessed how variability in spike timing (also called jitter^12,59^) is potentially shared between neurons. Trial-to-trial variability in this domain is much less explored than the spike count domain, but has been noted in V1 of anesthetized cats^9^ and in MT of awake monkeys^60^.

To directly address the question of correlated jitter within early visual populations, we used a measure inspired by spike count noise correlation: the Spearman correlation (ρ) of jitter across trials (henceforth notated ρ_jitter_). To quantify jitter for each cell, we use a Dynamic Time Warping (DTW) procedure adapted from^61^ (Figure 4A and see Methods). Briefly, a linear time warping procedure is used to fit a temporal shift for each trial that best aligns that trial with all of the other trials via a least-squares loss. When these shifts (Fig 4B) are correlated between a pair of cells across trials - that is, a pair of cells are “early” or “late” relative to themselves on similar trials - a pair of cells has a positive ρ_jitter_ (Fig 4C). Conversely, if one cell tends to “early” when the other is “late”, they have a negative ρ_jitter_. The linear DTW method straightforwardly extends to populations to estimate population shifts (ref ^61^, Fig 4A right side). We analyzed the correlation of the trial-to-trial shifts estimated from single cells with other single cells (ρ_jitter_) and single cells and variously constructed sub-populations of simultaneously recorded cells (ρ_global jitter_).

**Figure 4.**
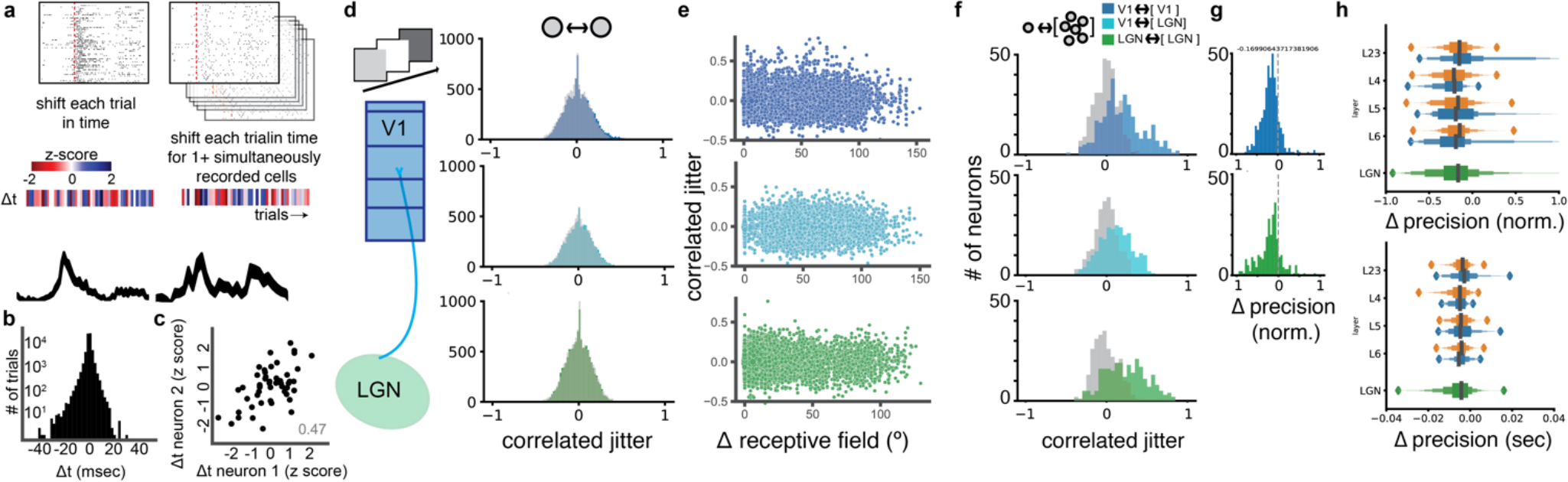
Correlated jitter across layers and areas in dLGN-V1 populations. **a.**, schematic illustrating the estimation of Δt on each trial for individual cells (left) and populations (right). For each cell (or population), the set of responses to an event or stimulus was fit via dynamic time warping, which assigns each trial a Δt to minimize the variance across trials (see Methods). Each Δt was zscored for measurement of correlated jitter (heat maps). **b.**, distribution of non z-scored Δt from dynamic time warping fits of each cell’s response to the flicker stimulus. **c.**, an example of a pair of single cells with positive correlated jitter. **d.**, correlated jitter (spike time noise correlation) for V1-V1 pairs (dark blue, top),_dLGN-V1 pairs (light blue, middle),and_dLGN-dLGN pairs (green, bottom). On each a null distribution from the same responses with trials shuffled for each cell is plotted in grey. **e.**, correlated jitter as a function of receptive field offset for V1-V1 pairs (dark blue, top),_dLGN-V1 pairs (light blue, middle),and dLGN-dLGN pairs (green, bottom). **f.**, correlated jitter (spike time noise correlation) for V1-V1 single neurons-populations (dark blue, top),_dLGN-V1 single neurons-populations (light blue, middle),and_dLGN-dLGN single neurons-populations (green, bottom). On each a null distribution from the same responses with trials shuffled for each cell is plotted in grey. **g.**, the normalized change in precision (first spike jitter), following correction of individual cells on a trial-by-trial basis with the Δt of the other cells in the same structure (V1 dark blue, dLGN green). **h.**, letter value box plots of the normalized change in precision (first spike jitter) in each layer, following correction of individual cells on a trial-by-trial basis with the Δt of the other cells in V1 (top) and dLGN (bottom).

The pairwise ρ_jitter_ across all V1 pairs (n = 33,207 pairs) was not significantly different than zero, with a mean of 0.004 ± 9e^−4^ (SEM, Fig 4D); unlike anesthetized cats^9^ this was not different from either zero (p < 1e^−16^, 1-sample T-test) or the null distribution of ρ_jitter_ expected from trial-shuffled data (p < 1e^−16^, Welch’s t-test). Within LGN, the ρ_jitter_ across all LGN pairs (n = 9929 pairs) was 0.002 ± 0.002 SEM (Fig 4D); this again was not significantly different from either zero (p < 1e^−16^, 1-sample t-test) or the null distribution of ρ_jitter_ expected from shuffled data (p < 1e^−16^, Welch’s t-test). In addition, ρ_jitter_ between pairs was not modulated by the retinotopic offset between pairs of cells, for any region (Fig 4E)

Despite not observing any pairwise correlated jitter, we nevertheless wondered if correlated jitter could contribute to population encoding. Weak pairwise correlations in rate variability have been shown to coexist within much stronger population correlations^31^. Given the noisiness in estimating a shift on each trial from individual cells, we hypothesized that the observed weak pairwise ρ_jitter_ metric had failed to capture shared shifts in latency among larger visual populations.

### Population correlated jitter: ρ_global jitter_

To assess the coupling of individual neuron jitter to the population, we adapted the pairwise ρ_jitter_. We estimated the Δt_population_ of population responses using the linear shift DTW method. We then defined the ρ_globaljitter_ for each single neuron as the Spearman correlation of the Δt_population_ with Δt of that neuron. The estimate of Δt_population_ never included the target single neuron, but could be made of variable populations of 1 to *n* cells, where n is the total simultaneously recorded cells; the case of *n*=1 is the same as pairwise correlated jitter described above.

Indeed, the ρ_global jitter_ of single cells within LGN and V1 to all other cells in their structure was positive. The ρ_global jitter_ of LGN cells to LGN populations had a mean of 0.38 ± 0.015 SEM (Fig 4B, green); this was significantly different from both zero (p < 1e^−16^, 1-sample t-test) and population control (p < 1e^−16^, Welch’s t-test). Similarly, the ρ_global jitter_ within V1 cells to V1 populations was non-zero, with a mean of 0.34 ± 0.007 SEM (Fig 4B, dark blue); this was significantly different from both zero (p < 1e^−16^, 1-sample t-test) and population control (p < 1e^−16^, Welch’s t-test). By computing the ρ_global jitter_ from individual cells in V1 with the Δt_population_ of LGN, we can estimate the amount of ρ_global jitter_ in V1 inherited from LGN. We found LGN population → V1 cell ρ_global jitter_ was non-zero, with a mean of 0.27 ± 0.02 SEM (Fig 4B, light blue); this was significantly different from both zero (p < 1e^−16^, 1-sample t-test) and population control (p < 1e^−16^, Welch’s t-test). By computing the ρ_global jitter_ from individual cells in LGN with the Δt_population_ of the V1, we can estimate the amount of ρ_global jitter_ in LGN imposed by feedback from V1. This V1 population → LGN cell ρ_global jitter_ was also non-zero, but significantly lower in magnitude, with a mean of 0.1 ± 0.01 SEM (Fig 4B, open light blue); this was significantly different from both zero (p < 1e^−16^, 1-sample t-test) and population control (p < 1e^−16^, Welch’s t-test).

The strength of ρ_global jitter_ in both LGN and V1 suggests that the apparent imprecision in single cell visual responses - when measured relative to stimulus time (Fig 1F), - may be an overestimation and responses are more precise within the context of the population encoding. To test this, we adjusted each cell’s response by the Δt_population_ on each trial and remeasured precision and reliability as we had previously. This resulted in a significant increase in single cell precision in LGN and in all V1 layers (Fig 4F,G; p < 1e^−23^, Welch’s t-test). The precision increased by 23% on average (13.2 + 3.4 msec) When adjusting by the total Δt_population_, there were no significant differences in precision increase across layers (Figure 4G).

In summary, trial-to-trial fluctuations in the *timing* of individual cells in V1 and LGN are strongly coupled to fluctuations in timing amongst the population. Single cells in V1 are coupled to LGN population jitter, but less strongly than to V1 population jitter, suggesting that some jitter correlation is inherited from “upstream” and some is added by local or feedback mechanisms in V1.

### Correlated variability subpopulations: layers and ensembles

The relationship of ρ_global jitter_ to the underlying functional architecture (i.e., retinotopy, V1 layers) could point to a mechanism through which shared jitter arises: shared inputs^62,63^, recurrent connectivity^64,65^, or network structure^66^. We first hypothesized, based on notions of a feedforward visual hierarchy^39,67^ with local recurrent connectivity^68,69^, that ρ_global jitter_ would be higher within groups of cells from the same layer. We tested this by measuring for each neuron the layer-specific ρ_global jitter_ (one neuron to all other neurons in a given layer or LGN). For both regular spiking and fast spiking cells, we found that ρ_global jitter_ was not significantly higher within-layer than across to some other layers, inconsistent with the hypothesis that ρ_global jitter_ is driven by within-layer recurrence (Fig 5A). In fact, LGN, which lacks local excitatory recurrence entirely, had the highest intrastructure ρ_global jitter_ (0.24, Fig 5A). The across-layer connections with the highest ρ_global jitter_ instead suggest correlated jitter is driven by specific feedforward connections: strongest coupling of V1 cells in the thalamorecipient zones to LGN fluctuations, stronger L5 coupling to L2/3 populations, and weaker L6 coupling to any V1 population. L2/3 seems to be the exception: V1 L2/3 single neuron ρ_global jitter_ is broadly high across V1 layers. This was true of each type of visual stimulus (Fig 5B). To quantitatively compare correlated jitter to feedforward connectivity and correlated spike count variability, we used a matrix similarity measure (Euclidean distance) to compare across-layer correlated jitter to across-layer connectivity (Fig 5C) and feedforward synaptic connectivity (as measured by paired *in vitro* recordings, in ref ^70^). This showed that the across-layer ρ_global jitter_ most resembled the spike count correlations in <10 msec bins (Fig 5C). The sole exception was the regular spiking cell response to natural movies, during which ρ_global jitter_ most resembled rate fluctuations on the seconds timescale. The across-layer correlated jitter was positively correlated across-layer connectivity (Fig 5D, r=0.42, p=0.14).

**Figure 5.**
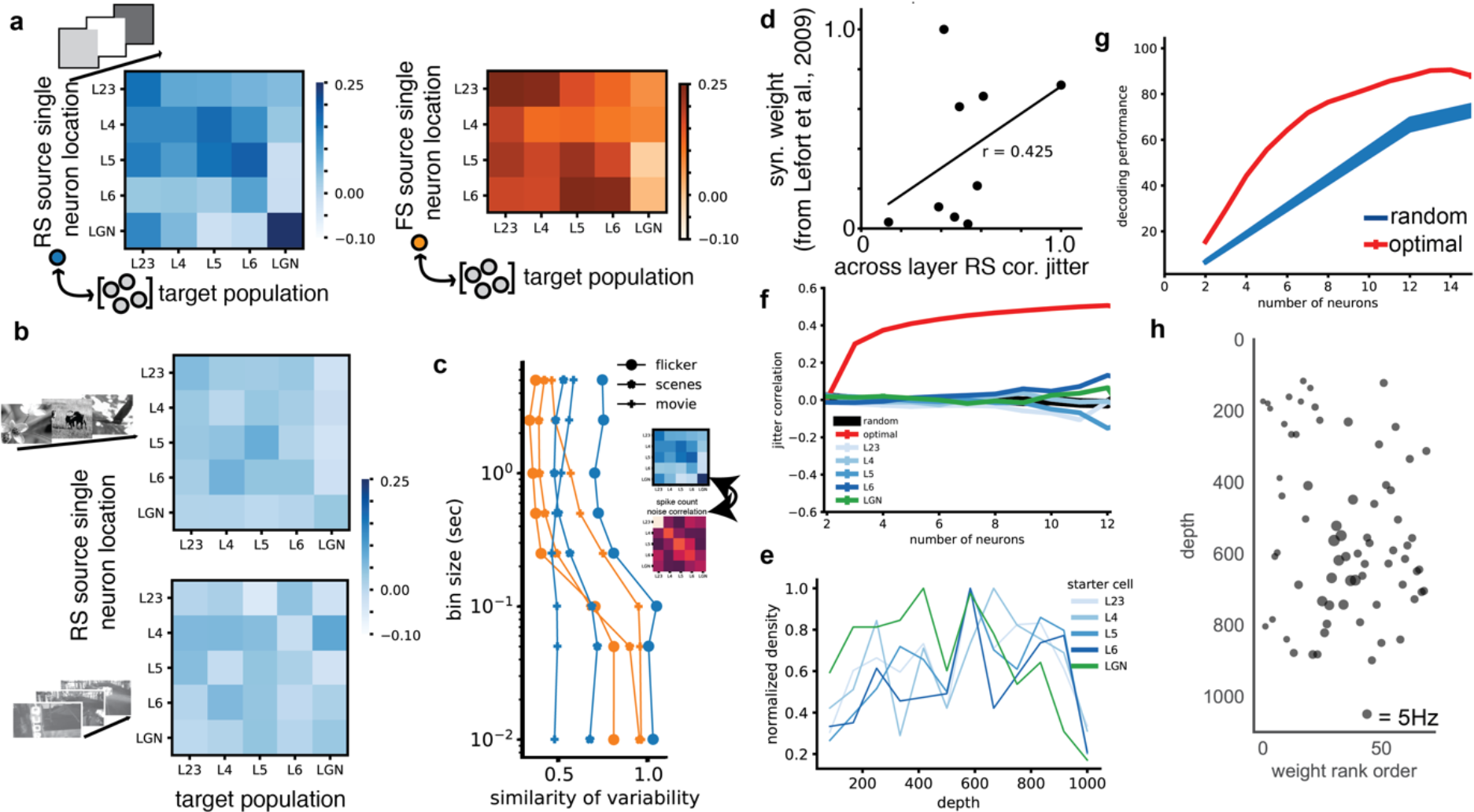
Correlated jitter laminar structure and interlaminar jitter ensembles. **a.**, the correlated jitter, ρ_global jitter_, of single cells in particular layers V1 or dLGN (left axis) with subpopulations of cells in particular layers V1 or dLGN (right axis). Single regular-spiking cells correlated subpopulations in blue, Single fast-spiking cells correlated subpopulations in orange, each measured with jitter in response to flicker stimuli. **b.**, single regular-spiking cells correlated measured with jitter in response to naturalistic image (top) and naturalistic movie stimuli (bottom). **c.**, the similarity of the single cell - V1 layer/dLGN jitter correlation to pairwise across layer/dLGN spike count correlation. Similarity is measured via the Euclidean distance (bottom axis) between the correlation matrices (inset), for correlated jitter of each stimulus type (symbols) and cell type (blue, regular-spiking; orange, fast-spiking), for spike count noise correlation at all bin size (left axis). **d.**, linear regression of regular-spiking cells subpopulation-jitter (from **a**), with the feedword connectivity strength (amplitude * probability) matrix from ref ^70^. **e.**, ρ_global jitter_ of the identified optimal ρ_global jitter_ sets (red), random sets of the same size (black), and random sets constrained to individual layers (blues) or dLGN (green). mean ± S.E.M. **f.**, the distribution across the cortical depth of the ρ_global jitter_ sets that originated in individual layers (blues) or dLGN (green). **g.**, performance of support vector machine (SVM) decoders trained and tested with cells with high correlated jitter (optimal ρ_global jitter_ sets), compared to random populations of the same size. **h.**, an example distribution of SVM weights from a decoder trained and tested with all simultaneous V1 cells, across the cortical depth. Symbol size indicates evoked firing rate, scale symbol = 5Hz.

The across-layer structure of ρ_global jitter_ also suggested that co-fluctuations in jitter could serve to link cell across layers. We wondered which sets of cells, within those simultaneously recorded, had the highest ρ_global jitter_. To find these sets, we began with, for each cell, the single other simultaneously recorded cell with the most correlated jitter; this other cell is the first member of the “optimal” ρ_global jitter_ set. We then iteratively added one cell at a time to the optimal set, correlating the jitter starter cell with the jitter of the new set. The next cell added to the set was the one that increased ρ_global jitter_ the most, and the optimal ρ_global jitter_ set grew until it adding more cells to ρ_global jitter_ set did not increase ρ_global jitter_.

## DISCUSSION

We found that neural populations in V1 and LGN have shared variability in spike count and spike timing, and that this variability enhances the visual stimulus information in population responses. Unmeasurable pairwise correlation in jitter manifests as strong population jitter correlations, revealing shared spike time variability. Further, we found that spike time variability is enhanced in subgroups of neurons that span layers. These inter-layer subpopulations are more informative population than layer-specific or random subpopulations. These measures of the structure of population activity suggests neural coding with precise spike times in visual cortex may be more tractable than previously estimated based on extrapolations from pairwise recordings.

### Synergy, redundancy and decoding

The amount of information in population responses has been a matter of considerable interest in sensory neuroscience. Although some reports indicated synergy^8,71^, a consensus formed around the idea that populations of neurons were redundant in the visual system^52,72–74^ as well as other sensory^75^ and motor cortex^76^. However, a recent report in awake, behaving primates reopened this issue, finding synergistic interactions between pairs of nearby neurons^51^ in visual cortex. Contrary to Nigam et al., we found that many pairs of nearby neurons in passively viewing mouse V1 are indeed redundant (Fig 2D), with only a few examples of strong synergy between nearby neurons. This discrepancy

Such “optimal” ρ_global jitter_ ensembles for each cell had, by definition, much higher ρ_global jitter_ than both random sets of cells of the same size and sets of cells confined to the same layer of the same size (Fig 5E). In contrast, random and layer-restricted sets had indistinguishable ρ_global jitter_ up to sets of 10 cells (Fig 5E, p > 0.05). Notably, the distributions of ρ_global jitter_ sets were similar across the depth of cortex for starter cells in all layers and LGN (Fig 5F, p > 0.05), indicating that “optimal” ρ_global jitter_ sets are not biased to the layer of the originating cell. For most (6/8) recordings, the observed “optimal” ρ_global jitter_ sets were a non-random subset of the possible sets from that recording (Kolmogorov-Smirnov test p > 0.05), with some cells appearing more and others less frequently than expected by chance. indicating that distinct correlated jitter defined groups of cells are present across layers within the population response.

We finally hypothesized that this across-area ρ_global jitter_ could be related to how the SVM utilized information from populations. Indeed, the SVM performed better than random cells when trained and tested on “optimal” ρ_global jitter_ sets, as identified by the above process (Fig 5F). This is consistent with what we had observed from decoding using the complete population responses (Fig 2) - top SVM weights were always distributed across layers (Fig 5G); these distributions were all also different than a random distribution (Kolmogorov-Smirnov test p > 0.05).

In summary, we analyzed the structure of variability in visual population responses, found certain cells are linked across layers and areas by fast co-fluctuations. We call these groups of cells contained more overall stimulus information. suggests that local synergy in V1 requires, in contrast to the passively viewing animals in this study, active engagement.

As our recordings yielded small populations, we were also able to further assess the additional information in population responses. In anesthetized macaque^52^, small populations are redundant up to 6 neurons; we also found redundancy/independence in small populations up to eight neurons. However, we then found a “sweet spot” of synergy with growing population size: redundancy did not increase with more neurons, but in fact tipped towards synergy for populations of 8-50 neurons. Beyond fifty, redundancy again prevailed. In addition to observing population synergy, we found that shuffling of trials reduced the decodable stimulus information when directly decoding population responses, another indication the concerted activity contains additional stimulus information.

### Shared variability: ρ_global jitter_

How might the concerted activity enhance stimulus information? The effects of spike count variability (“noise” correlation) on information has been extensively measured and theoretically discussed^77^. Although originally thought to be detrimental to population coding, these correlations can have either advantageous or detrimental effects on population decoding depending on the relationship between signal and noise correlation and the decoding mechanism. Spike count noise correlation in our data were correlated with signal correlations, consistent with a detrimental effect on population coding.

We also measured shared variability in timing independent of spike count, using a trial-to-trial dynamic time warping approach. While it appeared that this variability in jitter was independent from pairs of neurons (but see ^9^), by pooling multiple simultaneously recorded neurons we observed significant correlations in jitter. We suggest this form of correlated variability enhances population encoding. Using traditional linear readout decoding schemes, a non-zero ρ_jitter_ is most likely to be advantageous, especially when using time bins for counting less than the single cell jitter for that stimulus.

Our results indicate that relative timing can be of use in sensory coding, even in the absence of explicit representation of the global temporal reference frame^78^. This is intuitive: any neural readout of relative timing can only see the relative timing of its inputs, not anything about the external stimulus or even neural information beyond its inputs. Does actual neural decoding utilize the global coordination in jitter described here? We cannot strictly determine from the phenomenological description presented here if any neural decoder has an explicit representation of the shared temporal frame of reference, nor can we identify an explicit neural signature of the mechanisms that is causing these correlated fluctuations. However, population decoding need not have an explicit representation for such correlated jitter to matter: it follows from our observation of positive correlated jitter that temporal structure is preserved. And such temporal structure is known to affect both integration^17^ and propogation^79^. Because of this, a global jitter correlation necessarily impacts neural readout^80^.

### Comparison to other mammalian sensory systems

Multiple lines of evidence from audition and somatosensorial suggest that temporal variability can be shared in sensory responses, and serve to enhance the precision of population responses. The variability in spike timing in the visual response reported here, and indeed even the higher precision reported in anesthetized animals, is considerably higher than sensory responses in both the mouse auditory and somatosensory cortices^81^.

Why might visual responses be more temporally variable? While this has sometimes been attributed to a general increased excitability in visual circuits, leading to higher rates and more spontaneous firing, it may also be the case that experimentally-defined reference times (e.g., the stimulus presentation time) may more accurately track the ongoing neural common reference frame for non-visual sensory systems. The retina, with analog synapses and a range of dynamics, may introduce additional temporal variability when measuring central visual responses. In contrast, whisker tracking via motor efference signal of whisking^82^ may improve the estimation of stimulus time, and the subthalamic pathways are fast and reliable^83^. While no such efferent copy is monitored for auditory experiments, it may be the case that the cochlea is more accurate in transducing stimulus timing than the retina. Auditory cortex may have an explicit temporal reference frame^78^.

### Functional subcircuits

While we here measure the correlated jitter and found significant correlation of single cells to the population, we do not know what cells, from within V1 and LGN populations like those we recorded, are actually read out downstream of V1 (or LGN) by a neural decoding mechanism. Might functional subcircuits, known to exist in rodent V1^84^ have stronger latency variability correlation within these circuits? Indeed, we observed across-layer subcircuits that seemed to be linked by especially high correlated latency. How do these relate to underlying anatomy? Are these confederations connected to each other? Do they receive common input? Regardless of exact connectivity motif, might they have a higher degree of connectivity to each other in network topology? These questions remain to be tested via methods that can link structure and function, such as rabies tracing^85^ or electron microscopy^86^, in combination with high-density electrophysiology and optotagging. Whether or not these confederations that jitter together are related to anatomically defined subcircuits, they may form units for efficient downstream readout^87^.

### Candidate mechanisms

How could correlated jitter be utilized in the visual system? One hypothesis is that each response should generate synchronizing signals, for one or more reference frames. The onset transient, often overlooked or even removed in visual responses, has variable latency within V1 and could serve as such a synchronizing signal for separate or a single reference clock. Another possibility is eye movements, which can transiently change visual responses and could serve as an active means by which the top-down mechanisms that generate eye movement could modulate the temporal reference frame.

Several other candidate mechanisms present themselves. Such a global reference frame could be recovered from a running summation of the activity, similar to a global pooling; interneurons, specifically PV interneurons, are a good candidate for such a signal. Another alternative is the neural use of the observed oscillatory signals, such as gamma frequency (~40 Hz) oscillations. Gamma oscillations could be a reflection of the global clock signal or an active mechanism to enforce said clock through reverberant dynamics or even ephaptic coupling. In V1, individual units can exhibit varying levels of coherence with the cortical local field potential. Is ρ_jitter_ related it to spike-field coherence in V1? One argument against such a hypothesis is the lack of spike-field coherence in LGN. We hypothesize that spike field coherence in V1 may reflect the common temporal framework, but the field depends on the geometry of the underlying dipoles and is not necessary for the existence of a common temporal framework.

Finally, we cannot rule out that the observed correlated fluctuations are a result inherent only to passive-viewing, resulting from visual disengagement, shifts in spatial attention, or shifts general attention or other global state changes unrelated to visual coding. The observation of correlated jitter, and indeed synergistic population encoding in V1, will need to be verified in the context of active vision.

## METHODS

### Animals

All animal procedures were approved by the Allen Institute for Brain Science Institutional Animal Care and Use Committee (IACUC). Mice in this study were male C57BL\6J aged 60 – 182 days. For all mice, an initial headpost procedure was performed to attach a 10mm diameter circular opening aluminum head-fixation plate to the skull above the left hemisphere. During the procedure, the skin was resected above the skull above left hemisphere, the skull covered with MetaBond, and the opening sealed with Kwik-cast. The animal was allowed to recover for at least 7 days prior to habituation. Habituation to the recording apparatus extended over the period of one week, including head-fixation and presentation of each of the stimuli used during recording.

### Stimuli

Stimuli were presented using an immersive stimulation system, described in detail elsewhere^88^. In this study we used achromatic (for mouse) settings that produced a mean luminance of 5 cd/m^2^.

Stimuli were generally presented in the following sequence: a spatiotemporal dense noise stimulus, naturalistic images, spatially uniform flicker, a repeated naturalistic movie, and a second presentation of a spatiotemporal dense noise stimulus to measure continuity of response from beginning to end of the recording. The spatially uniform luminance flicker covered the entire visual field, and included step changes in luminance (+/− up to 80% of mean luminance) at 20Hz. Values of luminance transitions were chosen from a uniform distribution; the same sequence was used for all experiments. The total duration of the luminance flicker was 15 seconds and it was repeated 40-50 times. Naturalistic images (118 total images) were selected from the Berkeley Segmentation Dataset^38^ as described elsewhere^42^ and were scaled to cover 70° × 70°, beginning 10° into the left visual hemifield and extending mostly in to the right visual hemifield and beginning 15° below the horizon (parallel to mouse platform) and extending into the upper visual field. Images were presented at 10Hz with no inter-image stimulus. The naturalistic movie stimulus was scaled to cover the same area of the visual field, and consisted of repeated 30 second clip from the opening scene of the 1958 film *Touch of Evil*. This clip is a continuous tracking shot with no spatiotemporal discontinuities.

### In vivo electrophysiology

To record from small population across V1 and LGN we used one or two *Neuropixels* probes simultaneously. These are high-density electrophysiology probes with 276 – 384 contiguous channels on a single shank covering 2.76 – 3.84 mm. After head-fixation inside the stimulus environment, probes were inserted through an pre-drilled opening in the skull. Recordings were grounded to a screw inserted rostral to the visual areas, typically near contralateral motor cortex. Following insertion, the exposed skull was covered with 1-2% agar to attempt to improve recording stability and maintain brain health. Each probe was lowered V1, and for some recording where the geometry presented the opportunity, continuing through to subcortical structures. For all recordings, several channels were left outside of the brain but within the covering agar to facilitate data-driven probe localization (Figure S2 and below). The probe was allowed to settle for at least 15 minutes, but typically 30, before responses were recorded. While not all recording sessions included all stimuli, samples sizes for each stimulus are included throughout the paper.

Each recording included at least one probe inserted through V1 and a subset of probes also successfully spanned LGN and V1 on the same probe (Figure 1A,C). Following quality control, 1006 total single units were included from 13 recordings in 7 mice. A post-hoc assignment of both unit depth and, for visual cortex, cortical layer was made by taking the distance between the center of the waveform amplitude distribution along the shank and the channel estimated at just beyond the pial surface from a characteristic shift in alpha-band power (Figure S1F-H). Units were recorded from all cortical layers and across all putative sub-regions of dLGN, though both distributions were biased towards deeper regions of each structure (Figure 1C). All stimuli were presented using a modified DLP projection system (380 nm and 525 nm LEDs, mean 3.2 cd/m^2^) in a custom immersive visual environment^88^ covering all visual space sampled by the recorded neurons. All experiments included a white noise stimulus for characterization of linear spatiotemporal receptive fields; 54% of units had a significant linear receptive field (Fig S2). Both evoked and spontaneous firing rates were consistent with other reports for awake mouse V1 (V1 regular spiking: 13.8 +/− 0.32 Hz, V1 fast-spiking: 28.69 +/− 1.37 Hz, LGN: 17.02 +/− 0.63 Fig 2B, Durand et al 2016).

### Spike sorting and pre-processing

To recover the spike times from individual neurons recorded with the multi-electrode arrays, we used a workflow that included algorithmic and manual refinement steps. We began with spike time extraction and putative single unit isolation using Kilosort, and manually refined the results using phy (github.com/cortexlab/phy). During manual refinement, most decisions were merging of clusters that had been algorithmically split due to drift in amplitude on the timescale of the entire recording; in addition, the first spikes in a burst also needed to manually merged. Each unit was tagged as multi-unit (MUA) or single based on the waveform and shapes of neighboring waveforms and the middle few milliseconds of the spike time auto-correlograms. For each recording, the quality of isolation of each unit was quantified using several metrics: maximum signal-to-noise, purity of the auto-correlogram, and distance of the mean waveform from other units in both feature and waveform space. These metrics were combined into a single quality metric using a Fisher’s linear discriminant projection based on the manual labels. From each waveform, the spike duration, ratio of spike height to spike trough, and the slope of repolarization phase from trough to rest (Figure S1A-C). From these parameters a k means classification identified fast-spiking and regular-spiking neurons. Finally, the depth of each unit from the pial surface was calculated based on features of the neural data (Fig S1F-H). We calculated the power in a range of frequency bands for each channel for a short epoch. The distribution of several bands revealed features of the underlying structure; from the alpha-band (8-14 Hz, Figure S1H), we were able to identify the first channel above the pial surface. This channel served as the reference channel for measuring the depth of each unit. The center of each unit was calculated based on the mean waveform, weighted by the amplitude of the waveform on each channel.

### Data Analysis

Following pre-processing, all data were packaged into a pseudo-NWB^89^ format. These hdf5 files includes all spike and stimulus times sufficient to describe the experimental data collected. The responses of every cell to each stimulus type was quantified using established metrics in the field. Single cell latency variability, often called jitter, was measured as the variance in the first or median spike time. The trial-to-trial reproducibility of spike trains was computed using the normalized cosine similarity of each trial with all other trials. All analyses were performed using Python and available packages. Data and code are available at the following URL: github.com/danieljdenman/ V1TimingVariabiltyCorrelations

#### ρ_jitter_ and ρ_global jitter_ definition

To measure the correlations between cells across trials, we adapted a dynamic time warping method from ref 59. This procedure involves estimating, for responses to each event on each trial, a shift-only warping function that minimizes the difference between each trial and a template. The template was the trial averaged peri-event time histogram of a variable window following stimulus presentation” for flicker, this was a 100 msec window following luminance transition, for naturalistic scenes a 100 msec window following image transition, and for naturalistic movies a 100 msec window following identified response events^10^. The shift function was constrained such that the maximum shift on a given trial was ± 40 msec. So, for each individual cell and each event trial a −40 msec < Δt < 40 msec was estimated; to account for difference in jitter, this Δt was z-scored by the mean Δt for each cell. ρ_jitter_ was defined as the Spearman correlation of these z-scored Δt.

To estimate jitter in the simultaneous responses of more than on cell, we used the same process, except instead of fitting the response of one cell to its mean, we fit the shift-only warping model to *n*-dimensional mean, where *n* is the size of the population whose Δt_population_ being measured. This *n* varied from 2 cells to the total recorded, and was sometimes constrained to individual V1 layers, dLGN, or the subsets of the cells that had the highest ρ_global jitter_. Δt_population_ were also z scored. ρ_global jitter_ was defined as the Spearman correlation of a single cell’s z-scored Δt with these z-scored Δt_population_.

#### Information theoretic measurements

To measure the information in responses, we estimated total and noise entropy using the “direct method”^49^, following Reinagel and Reid, 2000. For each cell and for each visual stimulus, we estimated the total and noise entropies at several word lengths, from 2 to 64. From these word length estimates, an extrapolation to infinite word length the y-axis was made via a 2^nd^ order polynomial fit and used as the H_total_ and H_noise_ (Fig 2A). Information, in bits, is I = H_total_ − H_noise_. To estimate total entropy, we used a trial-to-trial shift method, where the response of a neuron on each trial was shifted by a random amount, with wrapping from the end of the trial to the beginning. These trains had the same total number of spikes and identical global statistics, but lacked the repeated trial-to-trial structure of actual responses. We also used generated rate-matched Poisson spike trains from the responses to repeated stimuli, and entropy estimates were similar (data not shown).

To estimate the synergy or redundancy in ensembles, we estimated I_sum_ and I_ensemble_ following ref. 53^53^. The I_sum_ is the sum of the stimulus information I from each neuron included in an ensemble, where I is calculated independently for each neuron as above. I_ensemble_ was calculated for the combined response of the neurons in an ensemble, where these combined responses consist of *k*\**m* vectors of the *k* neurons in the ensemble divided into *m* time bins. For the analyses shown here we used 1 msec bins. We repeated for ensembles of size 2 to *n*, where *n* is the number of neurons simultaneously recorded.

#### Support vector machine decoding

To analyze decodable information in the population across both population size and timescale, we trained and tested the performance of linear support vector machine (SVM) classifiers using the Python package scikit-learn. SVM classifiers were trained on 30 of the 118 images repeatedly presented in the naturalistic images stimulus; for each ensemble, we chose the 30 images that evoked the highest firing. We chose to use 30 images to adjust classifier performance to a reasonable range, as many images failed to evoke any responses in a given set of cells and therefore dragged overall performance down. We trained our SVM decoders using response vectors containing responses from 2 to *n* neurons, where *n* is the total number of neurons in a given recording. Each neuron’s response on each trial was divided in to *m* bins, with *m* determined by the bin size used for each instantiation of the decoder. The response on each trial thus consisted of an *n*\**m* length vector containing spike counts. For each Testing of decoder performance was done with cross-validation using 10% of the trials held out from the training set.

#### Statistics

Unless otherwise noted, we used Welch’s t-test for comparisons of distribution means with unequal variances and 1-sample t-test comparison to zero to test if distributions were different from a mean of zero. Where single p values were able to be encoded by a 16-bit floating point number they are reported, otherwise they are given as < 1e^−16^.

## ACKNOWLEDGEMENTS

We would like to thank Eric Shea-Brown, Alex Cayco-Gajic, Michael Buice, Stefan Mihalas, Daniel Millman, Saskia de Vries, Gabriel Koch Ocker, Xioaxuan Jia, and Brian Long for useful discussions in the preparation of this manuscript. We would also like to thank the Neurosurgery and Behavior team and Animal Care teams at the Allen Institute for Brain Science for assistance in animal surgery, care, and handling. We wish to thank the Allen Institute founders, Paul G Allen and Jody Allen, for their vision, encouragement and support.

## CONTRIBUTIONS

Conceptualization, DJD and RCR; Methodology, DJD; Investigation, DJD; Software, DJD; Writing – Original Draft, DJD and RCR; Writing – Review and Editing, DJD, and RCR.

## SUPPLEMENTAL FIGURES

**Supplemental Figure 1.**
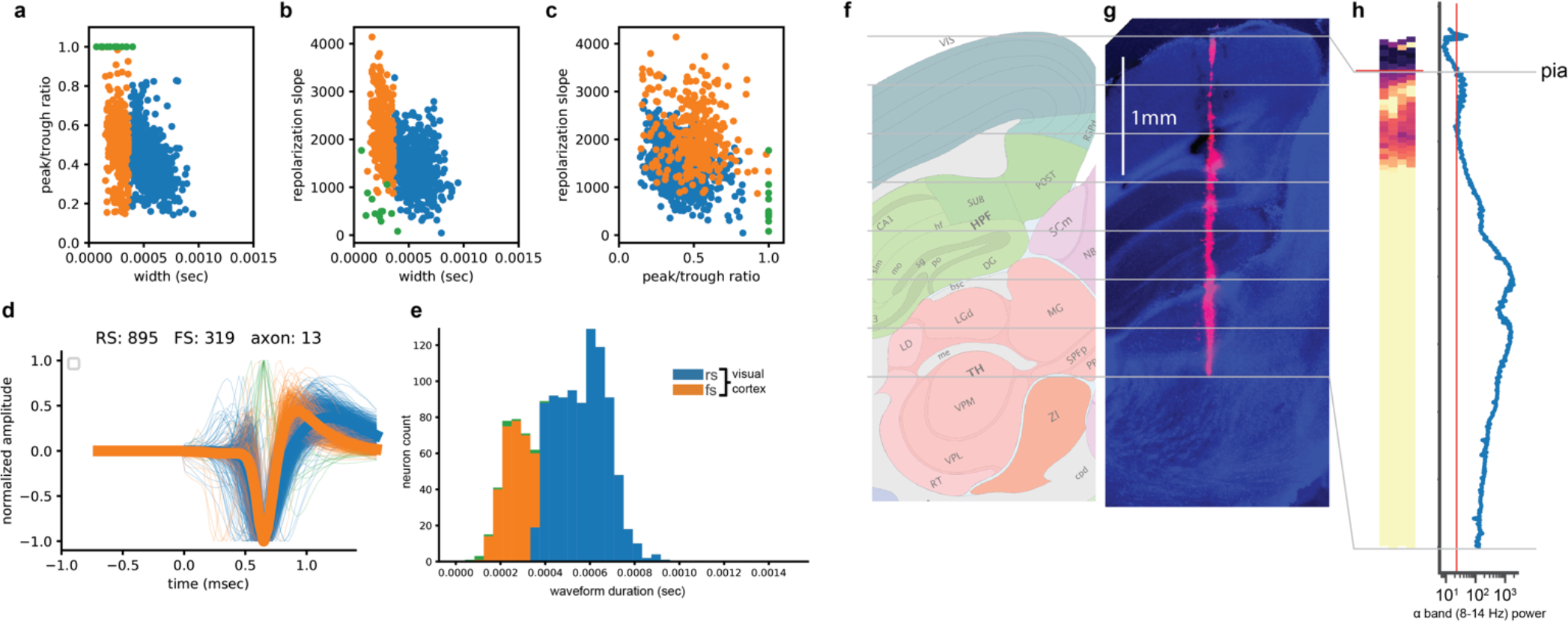
Assigning unit waveform type, depth, and layer. **a.-c.**, scatter plots of peak-trough amplitude ration, spike width (at half max), and repolarization slope for all units recorded from V1. Repolarization slope was calculated from a linear fit of the trough through the succeeding 246μsec. k-means (k=3) clustering was used to assign each unit an arbitrary label. Arbitrary labels were heuristically determined to be regular-spiking (RS), fast-spiking (FS), and positive-going groups. **d.**, individual waveforms (thin lines) and average waveforms for RS and FS groups. **e.**, histogram of spike widths. **f.**, Allen Reference Atlas coronal section corresponding to the electrode track in panel (**g**). **g.**, an identified electrode track imaged from 100μm coronal sections of fixed tissue; blue: DAPI, red: DiI. **h.**, the alpha-band power for each channel as a heatmap (left) and as a function of depth. Threshold crossings defined pia and white matter channels. Individual waveform depth, and subsequent layer identify, was measured based on the distance from the channel containing the peak waveform to this pia channel.

**Supplemental Figure 2.**
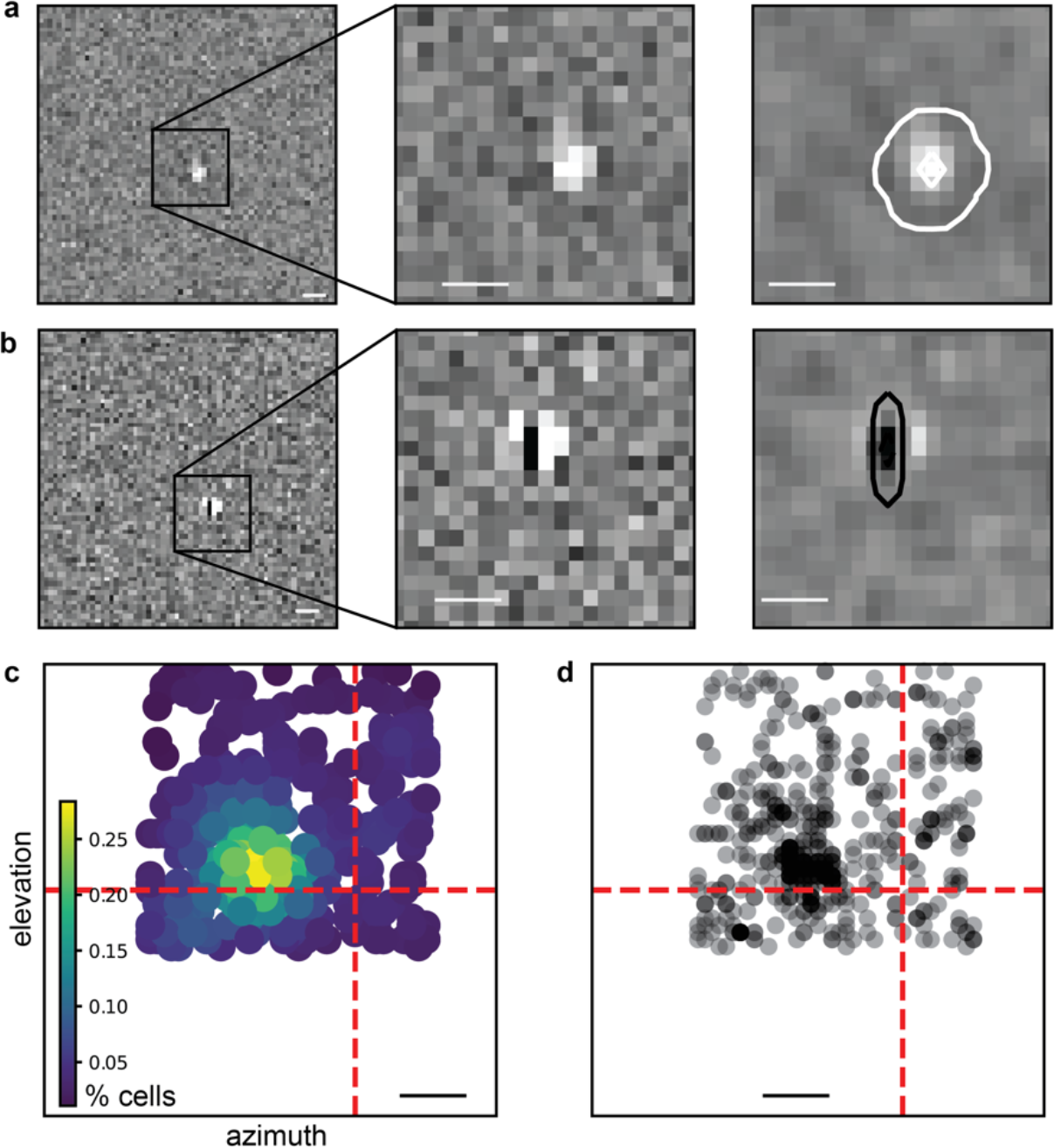
Receptive field mapping. **a.b.**, Example spatial receptive fields estimated from spike-triggered average of a ternary (black, white, and mean luminance grey) spatiotemporal noise stimulus. Leftmost panels show the complete 64×64 stimulus grid (3.4° square pixels), which covers the animals visual field. Middle panels show the spatial structure (top: center-surround and bottom: Gabor-like) **c.** the density of receptive fields across the stimulus display. **d.**, all individual receptive field locations. Scale bars in all panels are 17°.

**Supplemental Figure 3.**
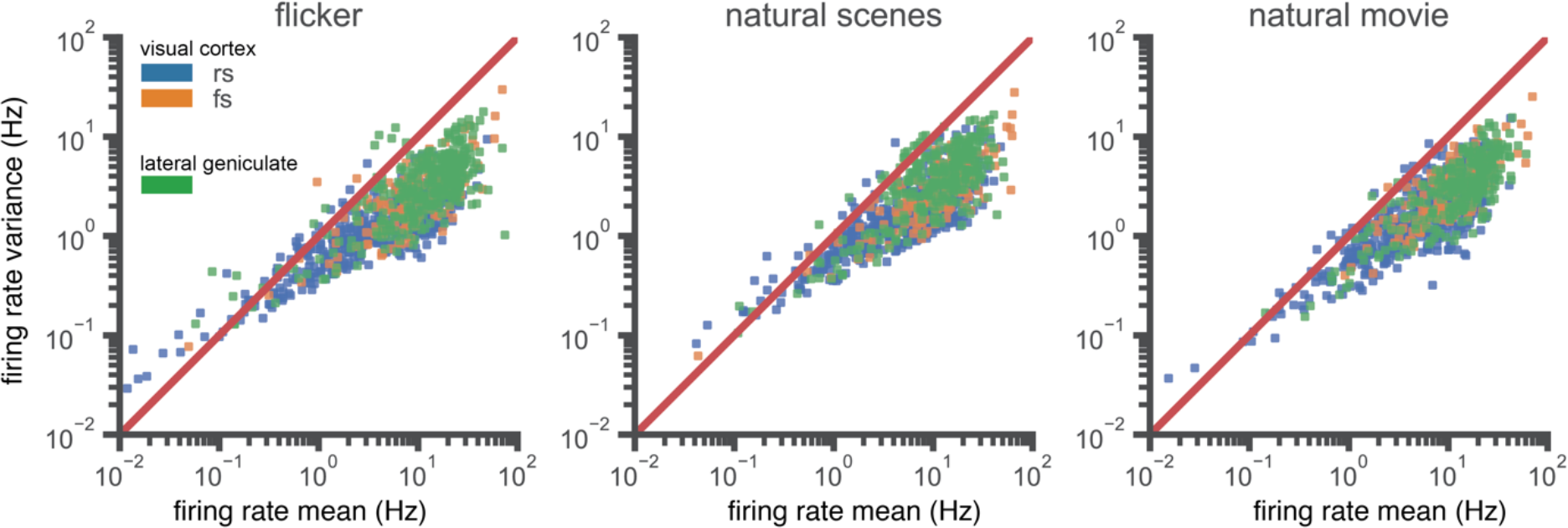
Sub-Poisson statistics for all stimulus types. Firing rate variance, for each of the three stimulus classes, as a function of the mean evoked firing rate. All axes are log-log.

**Supplemental Figure 4.**
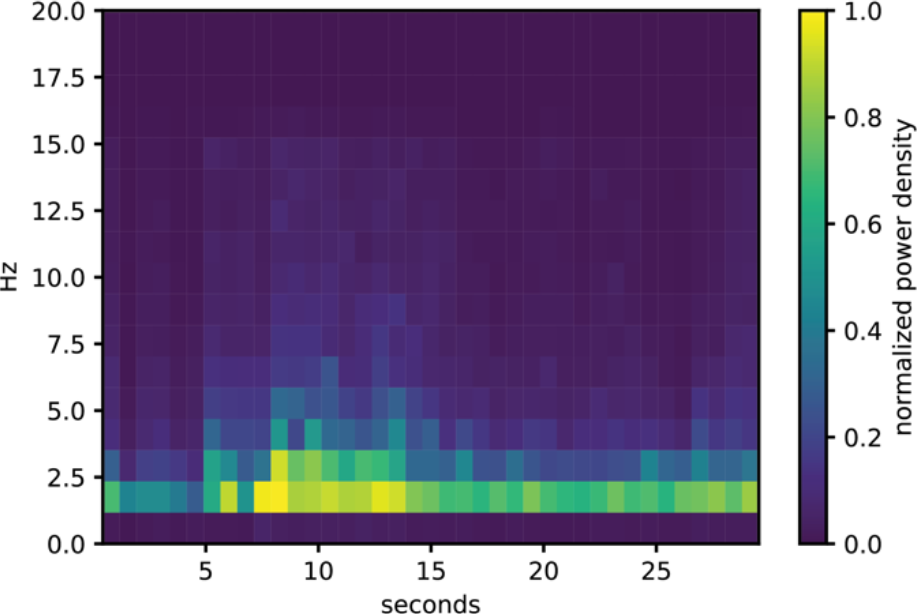
Specral analysis of naturalistic movie., Time-frequency spectrogram of the naturalistic movie clip (introduction to Touch of Evil), across the entire 30 second duration of the clip. The primary stimulus power is in the lower frequency bands, with a mean of 3.42Hz.

**Supplemental Figure 5.**
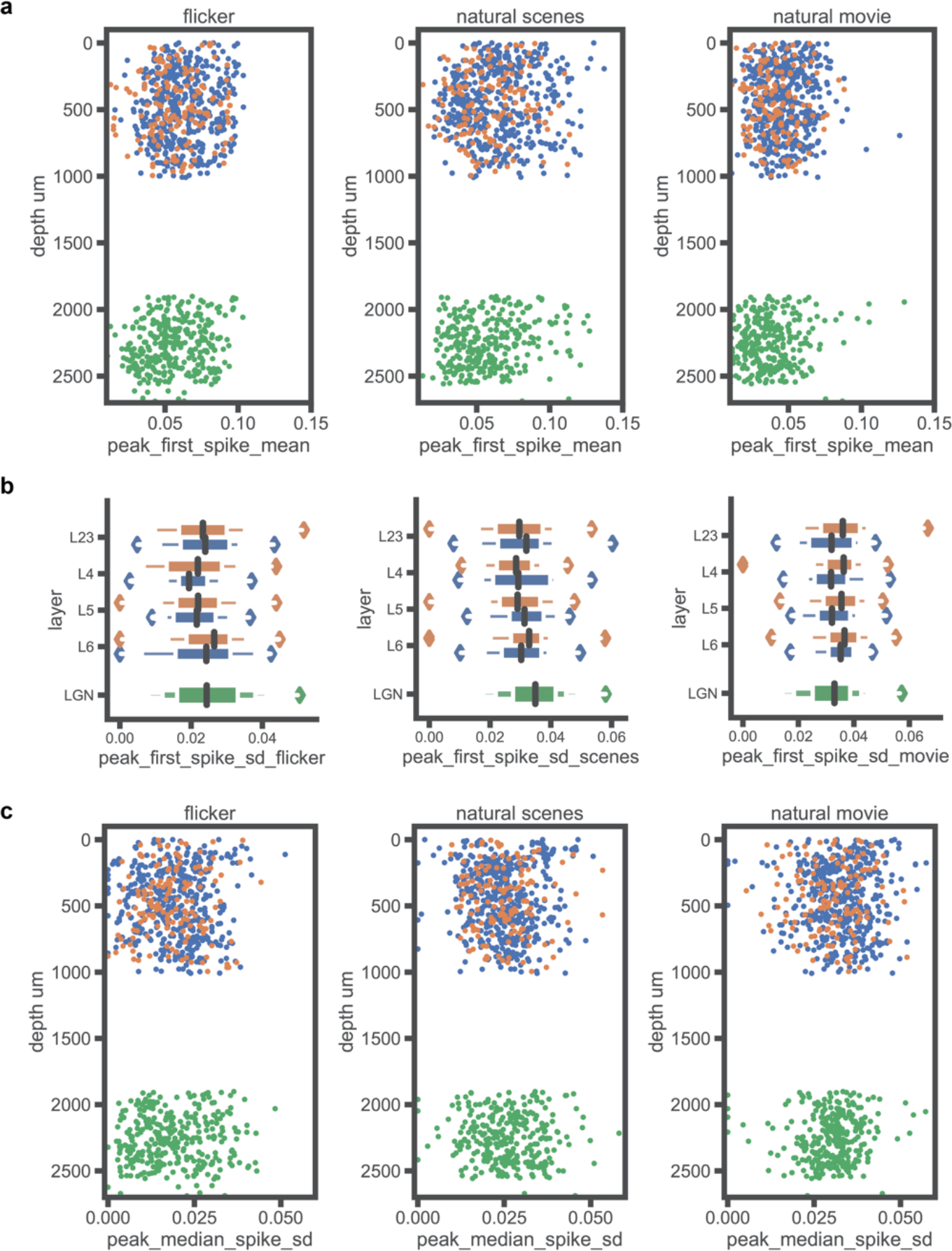
First spike latency and jitter. **a.**, absolute mean first spike latency for all cells as a function of depth, for each stimulus type. **b.**, letter value box plots, for each stimulus type, of the first spike jitter (standard deviation of first spike time on each trial, if there was a spike in the first 100 msec), broken down by layer for V1 and combined for dLGN. **c.**, as in **a**, but for median spike jitter.

**Supplemental Figure 6.**
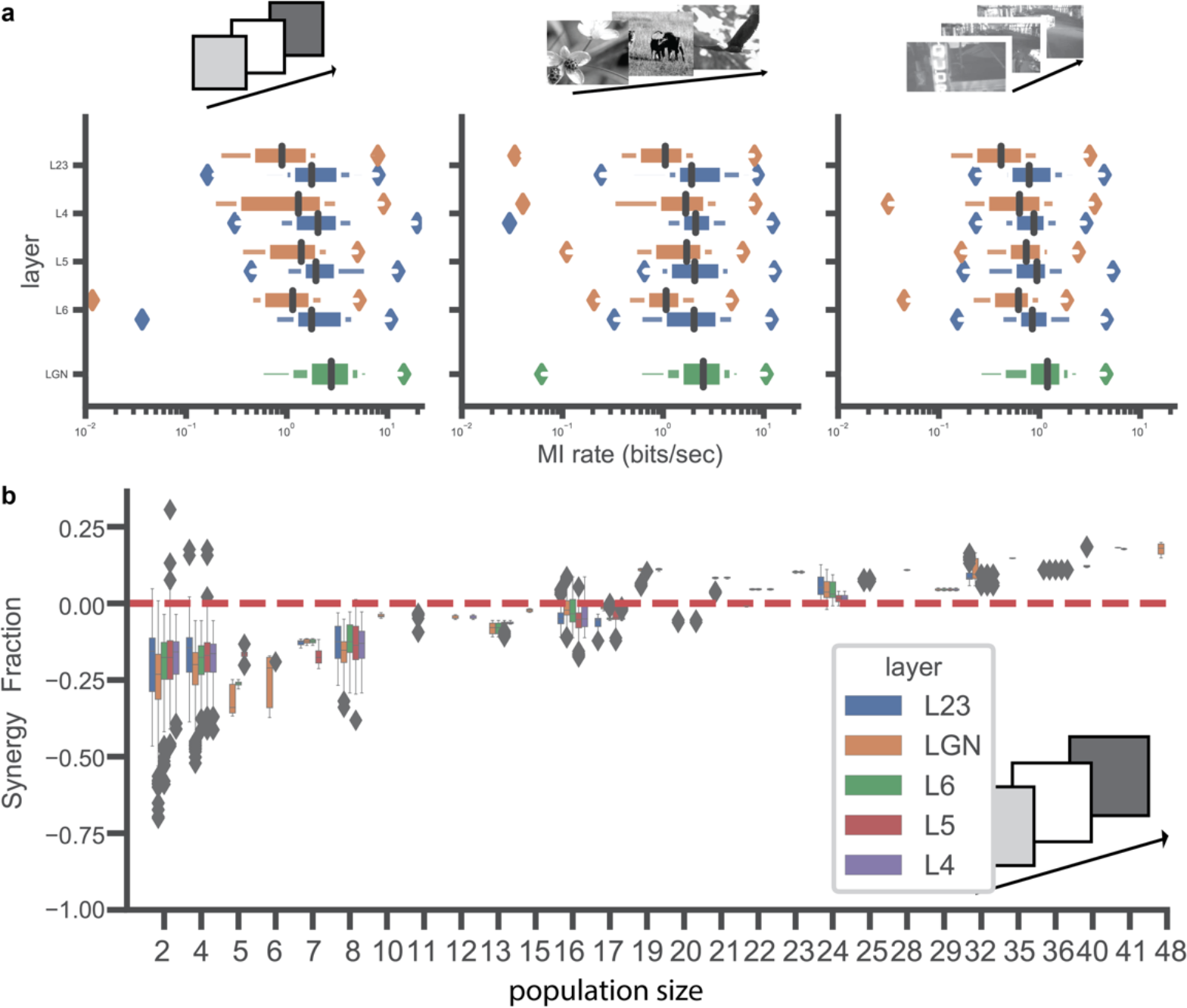
Receptive field mapping. **a** letter value box plots, for each stimulus type, of the mutual information rate, broken down by layer for V1 and combined for dLGN. **b.**, synergy fraction,(I_ensemble_ − I_sum_) / I_ensemble_ for flicker responses of each population of V1 layer or dLGN only sets as a function of number of cells in the set.

**Supplemental Figure 7.**
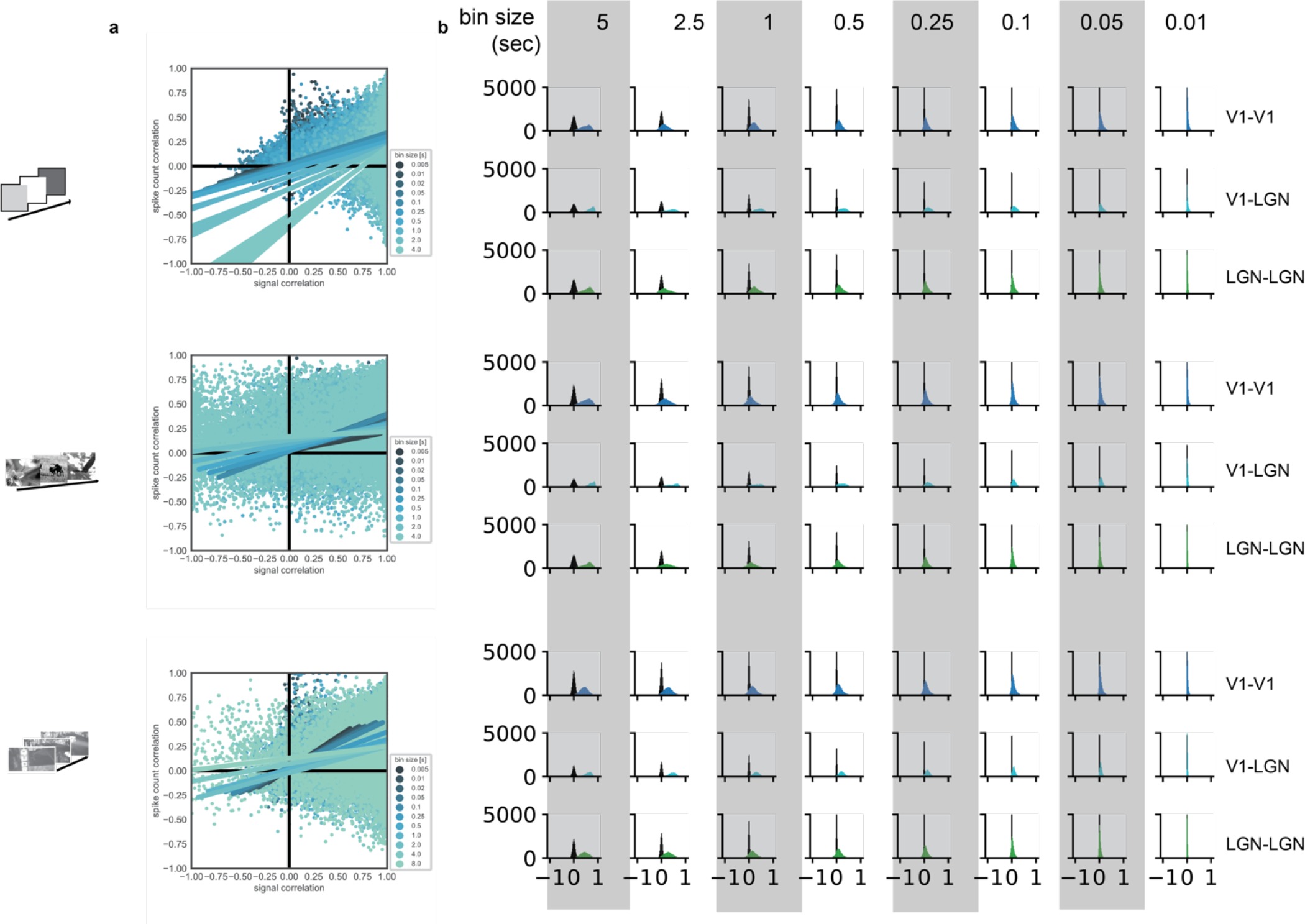
Signal and noise correlation. **a.**, Correlation of spike count signal correlation and noise correlation for all pairs at all timescales. Each stimulus type (rows) and timescales (colors) is individually fit with a linear regression. **b.** Spike count noise correlation for all pairs at all timescales. For each stimulus type (set of three rows) ans timescale (column) the V1-V1 pairs (dark blue, top),_dLGN-V1 pairs (light blue, middle),and_dLGN-dLGN pairs (green, bottom) are shown.

